# ATXN2 is a target of N-terminal proteolysis

**DOI:** 10.1101/2022.05.07.491020

**Authors:** Monika Chitre, Patrick Emery

## Abstract

Spinocerebellar ataxia 2 (SCA2) is a neurodegenerative disorder caused by the expansion of the poly-glutamine (polyQ) tract of Ataxin-2 (ATXN2). Other polyQ-containing proteins such as ATXN7 and huntingtin are associated with the development of neurodegenerative diseases when their N-terminal polyQ domains are expanded. Furthermore, they undergo proteolytic processing events that produce N-terminal fragments that include the polyQ stretch, which are implicated in pathogenesis. Interestingly, N-terminal ATXN2 fragments were reported in a brain extract from a SCA2 patient, but it is currently unknown whether an expanded polyQ domain contributes to ATXN2 proteolytic susceptibility. Here, we used transient expression in HEK293 cells to determine whether ATXN2 is a target for specific N-terminal proteolysis. We found that ATXN2 proteins with either normal or expanded polyQ stretches undergo proteolytic cleavage releasing an N-terminal polyQ-containing fragment. We identified a short amino acid sequence downstream of the polyQ domain that is necessary for N-terminal cleavage of full-length ATXN2 and sufficient to induce proteolysis of a heterologous protein. However, this sequence is not required for cleavage of a short ATXN2 isoform produced from an alternative start codon located just upstream of the CAG repeats encoding the polyQ domain. Our study extends our understanding of ATXN2 posttranslational regulation by revealing that this protein can be the target of specific proteolytic cleavage events releasing polyQ-containing products that are modulated by the N-terminal domain of ATXN2. N-terminal ATXN2 proteolysis of expanded polyQ domains might contribute to SCA2 pathology, as observed in other neurodegenerative disorders caused by polyQ domain expansion.

## Introduction

Spinal cerebellar ataxia (SCA2) is an autosomal-dominant, progressive, late-onset neurodegenerative disease that predominantly affects neurons in the cerebellum, brain stem, and spinal cord (1). Patients typically present with progressive cerebellar ataxia, ocular symptoms such as nystagmus or slow saccadic eye movements, and other neurological deficits such as peripheral neuropathy. SCA2 is associated with the expansion of the CAG-repeat region of the *ataxin 2* (*ATXN2)* gene and the resultant lengthened poly-glutamine (polyQ) tract that is located near the N-terminus of the ATXN2 protein (1–4). In the general population, the polyQ stretch usually contains 22 glutamines. Repeat lengths greater than 34 glutamines typically cause SCA2 (1–4). In addition, intermediate expansions of 27-33 glutamine repeats are associated with the development of amyotrophic lateral sclerosis (ALS) (5) and in some populations, Parkinson’s disease (PD) (6, 7).

The best characterized biochemical function of the ATXN2 protein is the regulation of mRNA stability and translation, including via miRNA silencing (8–14). ATXN2 is present in and regulates the formation of stress granules and P-bodies, which also control mRNA metabolism and translation (15–17). Many mRNA regulatory pathways have been implicated in the development of neurodegenerative diseases, with changes in polyQ length of specific proteins being a common feature (18). Indeed, polyQ lengthening is thought to influence protein-protein interactions, thus altering protein function and resulting in the formation of protein aggregates that might themselves be toxic.

To understand the role of ATXN2 in the development of neurodegenerative diseases, it is important to study its function as well as its regulation at both the gene transcription, mRNA and protein levels. Indeed, these mechanisms may prove to be promising therapeutic targets for reducing toxic protein loads. Transcriptionally, *ATXN2* expression is regulated by the transcription factor ETS1 (19). Interestingly, *ATXN2* pre-mRNA transcripts can produce multiple splice variants, but whether alternatively spliced *ATXN2* transcripts encode ATXN2 isoforms with differential activity or specific function is unclear (20). Furthermore, two potential start codons in the same translational frames are present in the 5’ end of the *ATXN2* gene and might thus add to isoform complexity (19). Post-translational regulation of ATXN2 has also been studied. For example, phosphorylation of ATXN2 by the cyclin-dependent kinase 5-p25 (Cdk5–p25) targets ATXN2 protein for degradation (21)

PolyQ proteins implicated in neurodegeneration can undergo an additional form of post-translational processing that could be targeted for therapeutic benefit. In the case of Huntingtin (Htt) (22) and ATXN7 (23), N-terminal proteolysis occurs near the N-terminal polyQ domain, thus releasing a polyQ containing fragment. Additionally, ATXN3 undergoes both N- and C-terminal proteolysis that produces N-terminal fragments as well as C-terminal fragments that contain its C-terminal polyQ domain (24, 25). Various proteases such as caspases, calpains, or matrix metalloproteinases target these polyQ proteins for cleavage (18, 23, 26). Strikingly, Huntington’s disease can be modeled in transgenic mice expressing the N-terminal fragment of mutant human Htt, suggesting that N-terminal cleavage products can be toxic (27). Moreover, inhibition of the protease that targets ATXN3 (25) or expression of cleavage-resistant forms of Htt (28) or ATXN7 (23) delays disease progression in animal models.

Despite the presence of an N-terminal polyQ domain in ATXN2, it has yet to be determined whether ATXN2 might also be the target of specific N-terminal proteolysis. Interestingly, an early study using brain extract from a SCA2 patient detected polyQ containing ATXN2 fragments, implicating N-terminal proteolysis of ATXN2 (29). Here, we present evidence that ATXN2 indeed can undergo specific N-terminal proteolytic cleavage, irrespective of polyQ length. This was the case for both long and short ATXN2 isoforms produced from alternative ATG start codons. We precisely map the cleavage site for the long isoform to a short and highly conserved motif located just C-terminal of the polyQ domain.

## Results

### ATXN2 undergoes N-terminal proteolysis in HEK293E cells

To determine whether ATXN2 can undergo N-terminal proteolytic cleavage, and whether an N-terminal polyQ-containing product might accumulate, we expressed full length ATXN2 proteins in HEK293 cells, using constructs that included the most 5’ of the two known potential ATG start codons (19) and an N-terminal HA tag, since there is currently no available N-terminally-directed anti-ATXN2 antibody (Fig. 1A). We expressed both ATXN2 with a normal polyQ domain of 22 amino acids (ATG1-HA-ATXN2-22Q), and ATXN2 with a pathogenic repeat of 39 glutamines (ATG1-HA-ATXN2-39Q). Extracts from these transfected cells were immunoblotted using both C-terminally-directed ATXN2 (C-ATXN2) and N-terminally-directed HA antibodies (Fig. 1B). The C-ATXN2 antibody detected two bands that could correspond to the full-length protein: 1) a band migrating with an apparent molecular weight of ~145 kDa that comigrated with endogenous ATXN2 detected in untransfected cells, and 2) an unexpected band migrating at ~180 kDa). Interestingly, the HA antibody detected the ~180 kDa isoform, but not the ~145 kDa band (Fig. 1B). Furthermore, there was a subtle, but reproducible difference in migration between the slower migrating bands observed with wild-type (22Q) and pathogenic (39Q) polyQ domains, while the migration of the ~145 kDa band was insensitive to polyQ length. These results suggested that the ~180 kDa band contains both the N-terminal and polyQ domains, while the ~145 kDa band does not contain the N terminus, possibly due to proteolytic cleavage. Consistent with this premise, we detected small HA-immunoreactive bands migrating at ~27 and 30 kDa in extracts from cells expressing ATG1-HA-ATXN2-22Q and ATG1-HA-ATXN2-39Q, respectively (Fig. 1B), suggesting that both normal and mutant ATXN2 proteins are subject to proteolysis. The difference in migration between the two fragments when probing for HA suggested that these fragments contain the polyQ domain and migrate differently due to the differential length of the polyQ domain. We noted that an additional band sensitive to polyQ length was consistently detected at ~65-70 kDa with the HA antibody (Fig S1A). A ~115 kDa band was also observed with the C-terminal ATXN2 antibody in lysates expressing both HA-tagged or untagged ATG1-ATXN2 lysates, and the molecular weight of this band did not vary in response to changes in polyQ length (Fig S1A, B). Based on their apparent sizes, these two bands could be the product of a second proteolytic cleavage event. The signal strength of the ~115 kDa band varied across experiments as opposed to the more stable ~180 and 145 kDa bands. We therefore focused this study on the proteolysis event that generates the N-terminal ~27 kDa and C-terminal ~145 kDa fragments.

**Figure 1.**
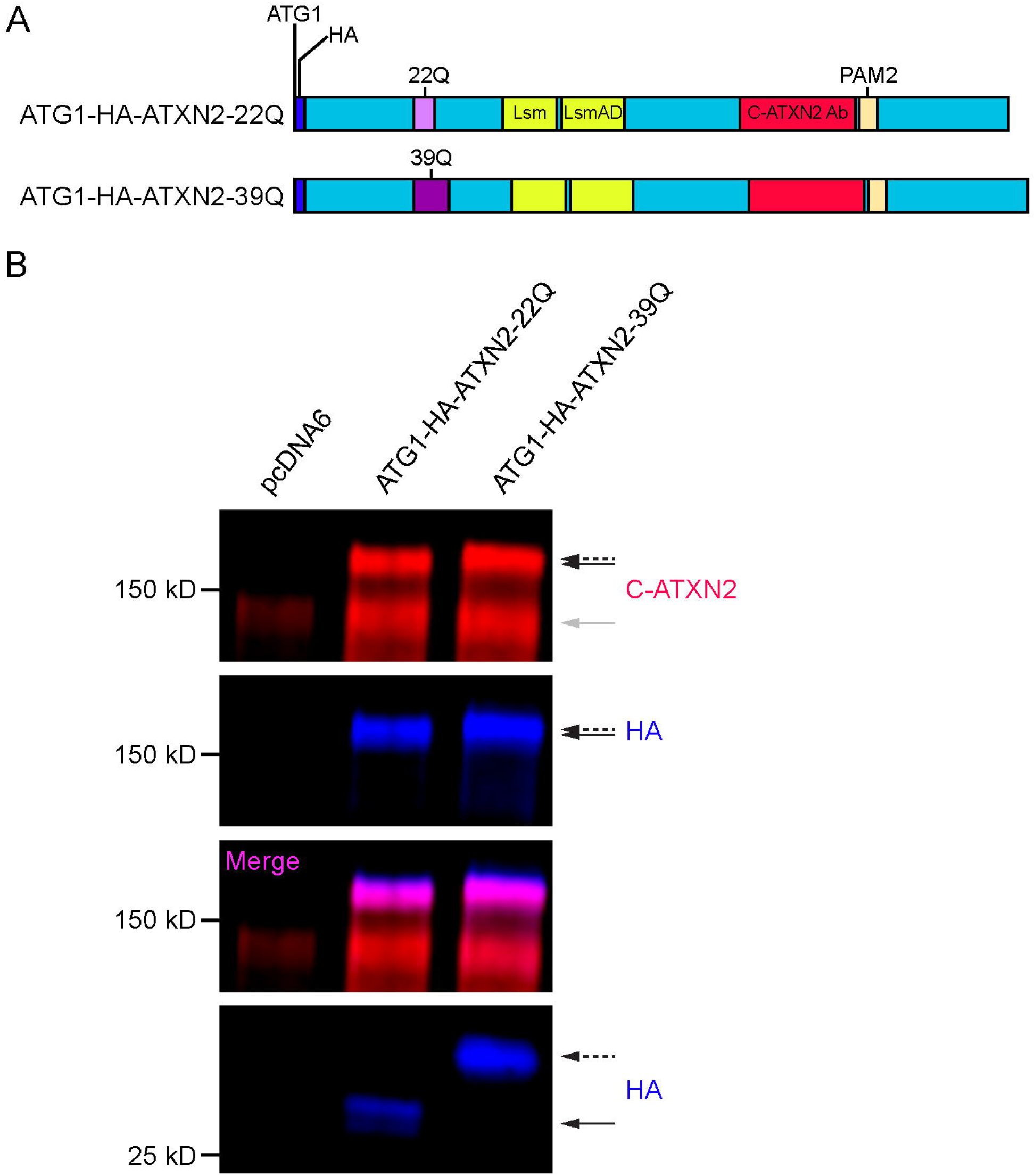
ATXN2 is a target of N-terminal proteolysis when transiently overexpressed in HEK293E cells. A. Schematic of HA-tagged full-length (1313 amino acid long) ATXN2 protein, starting at its 1^st^ ATG start codon (gray). An N-terminal tag (dark blue) was inserted immediately after the 1^st^ ATG for both the WT (ATG1-HA-ATXN2-22Q, light purple) and mutant (ATG1-HA-ATXN2-39Q, dark purple) forms of ATXN2 in a pcDNA6 vector backbone with a CMV promoter. B. Western blot of control HEK293E lysates transfected with empty vector (pcDNA6) vs. WT (ATG1-HA-ATXN2-22Q) and mutant (ATG1-HA-ATXN2-39Q) ATXN2 constructs. Probing for the C-terminus of ATXN2 (C-ATXN2, red) yielded an expected ~145 kDa band (gray arrow) in addition to a ~180 kDa (white asterisk) band in both WT and mutant ATXN2 lysates. Probing for the N-terminus of ATXN2 (HA, blue) only detected the ~180 kDa band in WT (black arrow, solid tail) and mutant (black arrow, dashed tail) ATXN2 lysates. The Licor imaging system allows for probing of two separate antibodies in the same lysate via conjugated fluorescent secondary antibodies that are visualized in separate channels in a similar fashion to immunofluorescence microscopy. The merged image of the two channels (magenta) shows that only the ~180 kDa bands (black arrows) can be simultaneously detected by both the N-terminal HA tag (blue) and C-terminal ATXN2 antibody, clearly indicating that this band is a full-length isoform of WT or mutant ATXN2. We also detected HA+ N-terminal fragments that migrate at ~27 kDa for WT ATXN2 (black arrow, solid tail), while the mutant ATXN2 lysates produce an HA+ N-terminal fragment that migrates at ~30 kDa (black arrow, dashed tail), suggesting that the N-terminal fragments include the polyQ domain. A Western blot representative of four independent experiments is shown.

To determine whether the ~27 kDa N-terminal fragment and the ~180 kDa ATXN2 isoform indeed contain the polyQ repeat, we used an anti-polyQ antibody (1C2) (30) that detects extended polyQ domains, but not shorter, wild-type polyQ repeats (31). 1C2 detected a polyQ stretch in both the small HA-positive N-terminal fragment and the ~180 kDa isoform from ATG1-HA-ATXN2-39Q lysates, whereas 1C2 did not detect a polyQ stretch in the N-terminal fragment from ATG1-HA-ATXN2-22Q lysate, as expected (Fig. 2A).

**Figure 2.**
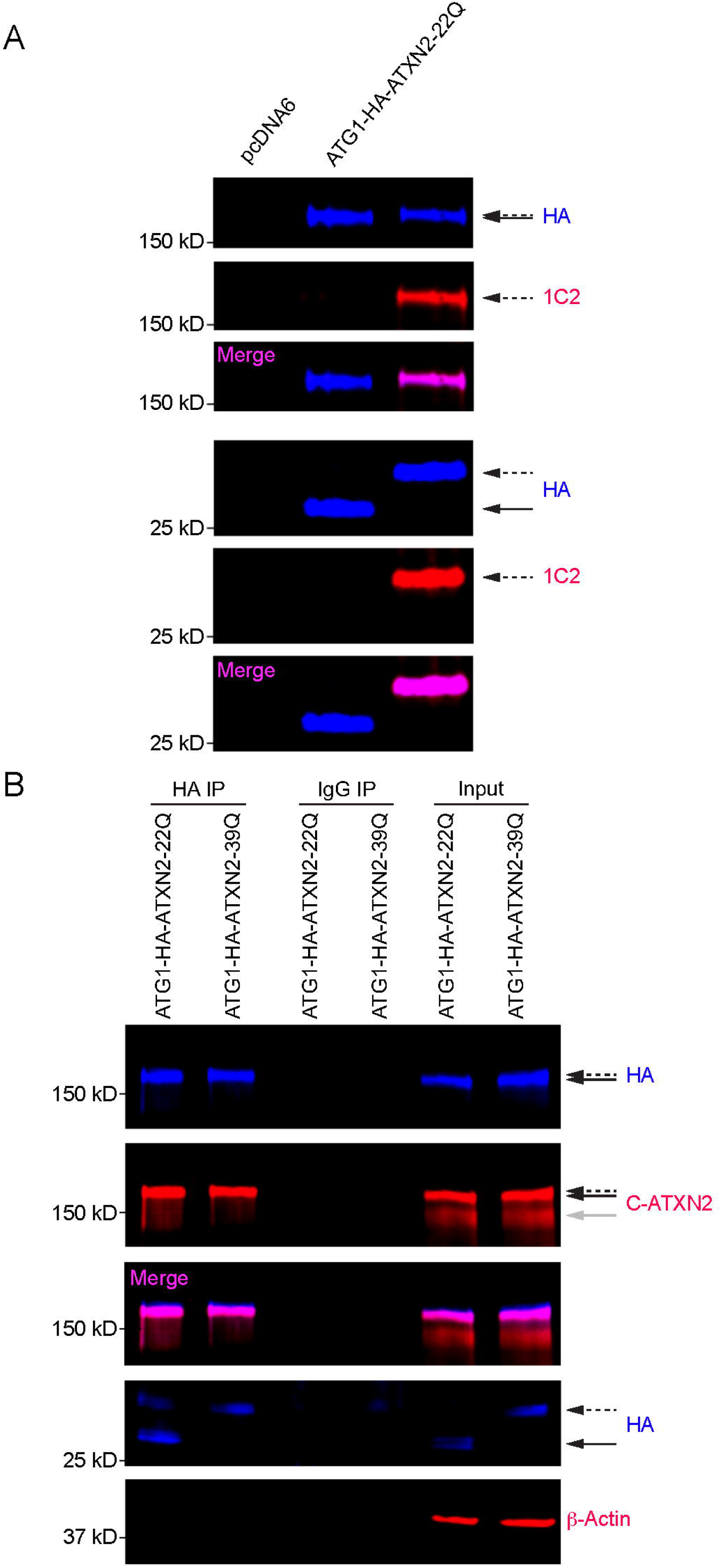
N-terminal proteolysis produces a N-terminal fragment that contains the polyQ domain. **A**. HEK293E lysates probed for the polyQ domain with the HA (blue) and 1C2 antibody (red), the latter of which recognizes only the expanded polyQ domain. As expected, only the mutant ATXN2 lysates showed a signal with 1C2: a band at ~180 kDa (black arrow, dashed tail) and a N-terminal fragment at ~30 kDa (black arrow, dashed tail) were detected. The polyQ signal overlays with HA signal when merging the two channels in the mutant ATXN2 lysates (magenta). A Western blot representative of three independent experiments is shown. **B**. ATXN2 was immunoprecipitated with HA antibody from HEK293E cells transfected with either 22Q or 39Q ATXN2. Rabbit IgG was used as an immunoprecipitation control. When probing for the N-terminus of ATXN2, only the full-length (180 kDa) protein and N-terminal cleavage fragments were detectable following immunoprecipitation with HA antibody. A Western blot representative of three independent experiments is shown. Image brightness and contrast were adjusted for better visualization of the small fragment in panel B.

To investigate whether the N-terminal and C-terminal fragments of ATXN2 remain associated after cleavage, we expressed ATG1-HA-ATXN2-22Q or ATG1-HA-ATXN2-39Q and performed immunoprecipitation with anti-HA antibodies. Full-length ATXN2 and the N-terminal fragment were efficiently immunoprecipitated with the HA antibody, but not the ~145 kDa C-terminal fragment (Fig. 2B). These results suggested that upon proteolytic cleavage, the N-terminal fragment does not remain associated with the rest of the protein, and further validated that the ~145 kDa band lacks an N-terminus.

In conclusion, when overexpressed, ATXN2 undergoes N-terminal proteolytic cleavage. The slower migrating isoform is the full-length protein, while the ~145 kDa protein isoform is a cleavage product containing all known functional protein domains but lacking the polyQ domain. Finally, a small N-terminal fragment containing the polyQ domain accumulates.

### A 17 amino acid domain C-terminal of the polyQ domain is necessary and sufficient for N-terminal cleavage of ATXN2

We next sought to understand how ATXN2 undergoes proteolytic cleavage and leveraged a deletion mapping strategy to identify the key protein domain(s) promoting its cleavage. Since the banding patterns described above suggested that the cleavage occurred downstream of the polyQ domain, we first generated a construct encoding a protein fusion comprising 395 N-terminal amino acids of ATXN2 and EGFP, with a short linker sequence separating the two moieties (Fig. 3A). This fusion protein was proteolytically processed in a similar way as full-length ATXN2, producing the same N-terminal cleavage fragment seen in ATG1-HA-ATXN2-22Q lysates. This indicated that the C-terminal region of ATXN2 is not required for N-terminal cleavage to occur (Fig. 3B).

**Figure 3.**
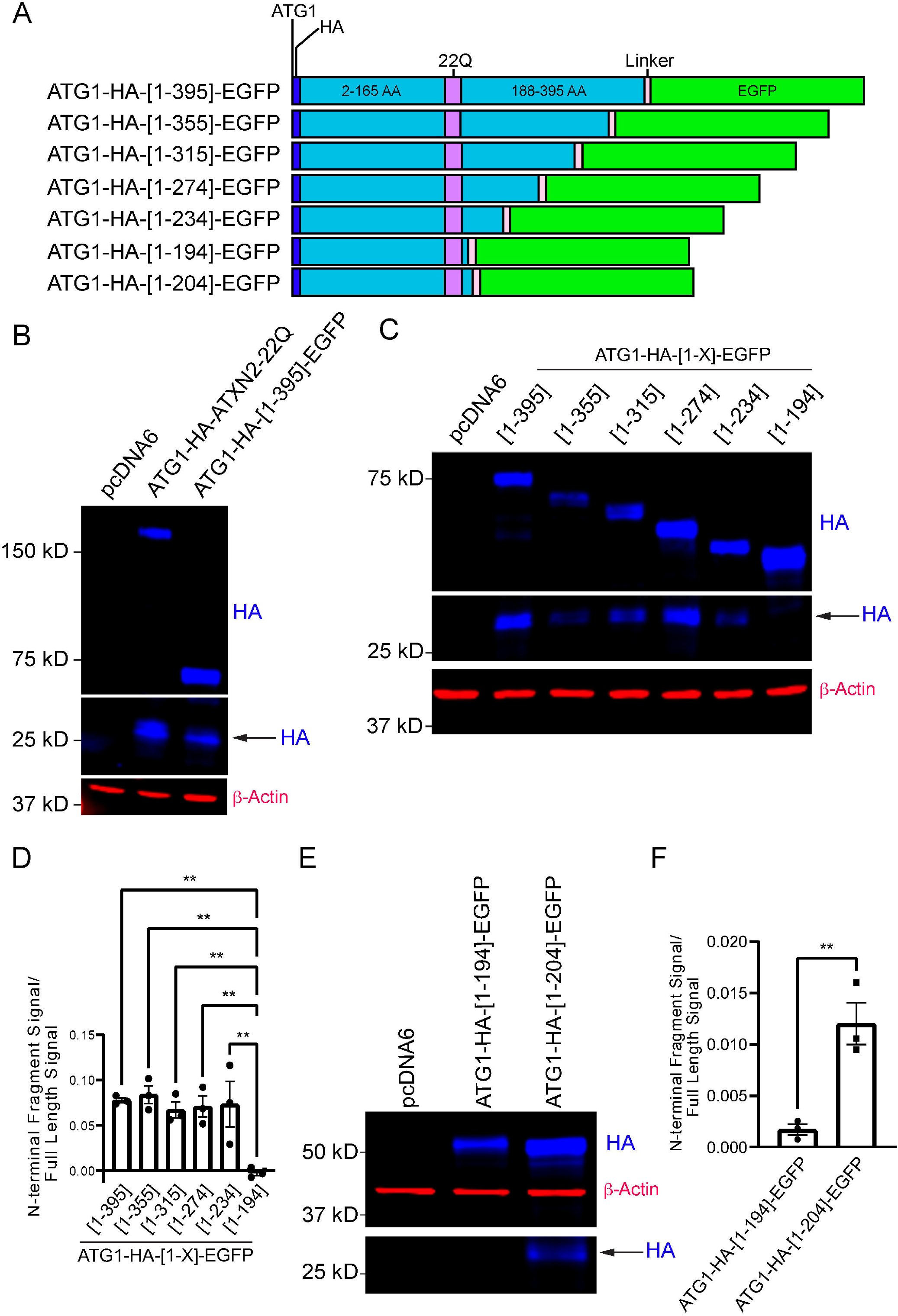
N-terminal cleavage of ATXN2 is abolished after deletion of amino acids adjacent to the polyQ domain. **A**. Schematics of proteins expressed in HEK293 cells that comprise ATXN2 N-terminal fragments of various lengths fused to a flexible amino acid linker sequence (black and white diagonal stripe) and a C-terminal EGFP reporter (green) **B**. HEK293E cells transfected with ATG1-HA-[1-395]-EGFP produced a cleavage fragment that migrates at the same molecular weight as full-length ATG1-HA-ATXN2-22Q (blue)**. A** Western blot representative of three independent experiments is shown. **C**. HEK293E cells transfected with N-terminal fragment constructs all produced N-terminal cleavage fragments detectable via HA (blue) except for the shortest construct (black arrow), indicating that a region immediately following the polyQ domain controls N-terminal cleavage of ATXN2. A Western blot representative of three independent experiments is shown. **D.** Quantification of N-terminal fragment cleavage efficiency (signal intensity of N-terminal fragment divided by that of full-length protein). Statistical analysis performed with one-way ANOVA with Dunnett’s multiple comparisons test. **P < 0.01 **E**. Western blot probing for HA (blue) shows that reintroducing 10 amino acids to the sequence downstream of the polyQ domain restores cleavage and produces an N-terminal fragment at ~27 kDa (black arrow). A Western blot representative of three independent experiments is shown. **F.** Quantification of N-terminal cleavage efficiency. Statistical analysis performed with unpaired T-test. **P < 0.01 Image brightness and contrast were adjusted for better visualization of the small fragment in both panel C and E, as cleavage efficiency was quite low in these experiments.

The N-terminal domain was then progressively deleted further in the C to N direction in increments of 40-41 amino acids (Fig. 3A). Cleavage and generation of an N-terminal fragment was observed with all fusion proteins, except with the fusion that included only the first 194 amino acids of ATXN2 (i.e. it contained only 7 amino acids after the polyQ, Fig. 3C). Thus, a sequence immediately following the polyQ domain is needed for N-terminal proteolysis (Fig. 3D). To refine mapping of the cleavage-promoting site, 10 amino acids were added back to produce an ATXN2-Nt-EGFP fusion that included the 17 amino acids immediately C-terminal of the polyQ domain (Fig. 3A). Reintroducing these 10 amino acids to the N-terminal domain restored N-terminal proteolysis of the fusion protein, which indicated that the 17 amino acids following the polyQ are critical for ATXN2 cleavage (Fig. 3F, G).

The above results point to a cleavage motif or perhaps a protease recognition site adjacent and downstream of the polyQ domain. To determine whether this short domain is necessary for cleavage, we internally deleted the 17 amino acids immediately downstream of the polyQ domain from the ATG1-HA-ATXN2-22Q protein (Fig. 4A). This deletion completely abolished cleavage, demonstrating the necessity of this short sequence for N-terminal proteolysis of full-length ATXN2 (Fig. 4B, C). Interestingly, elements of this unique sequence are perfectly conserved in mammals from mice to humans regardless of whether ATXN2 contains a polyQ domain or not. The amino acid sequences flanking the conserved sequences are also repetitive in nature and similar in terms of amino acid content, suggesting that this sequence may play an important role in ATXN2 function or metabolism (Fig. 4D).

**Figure 4.**
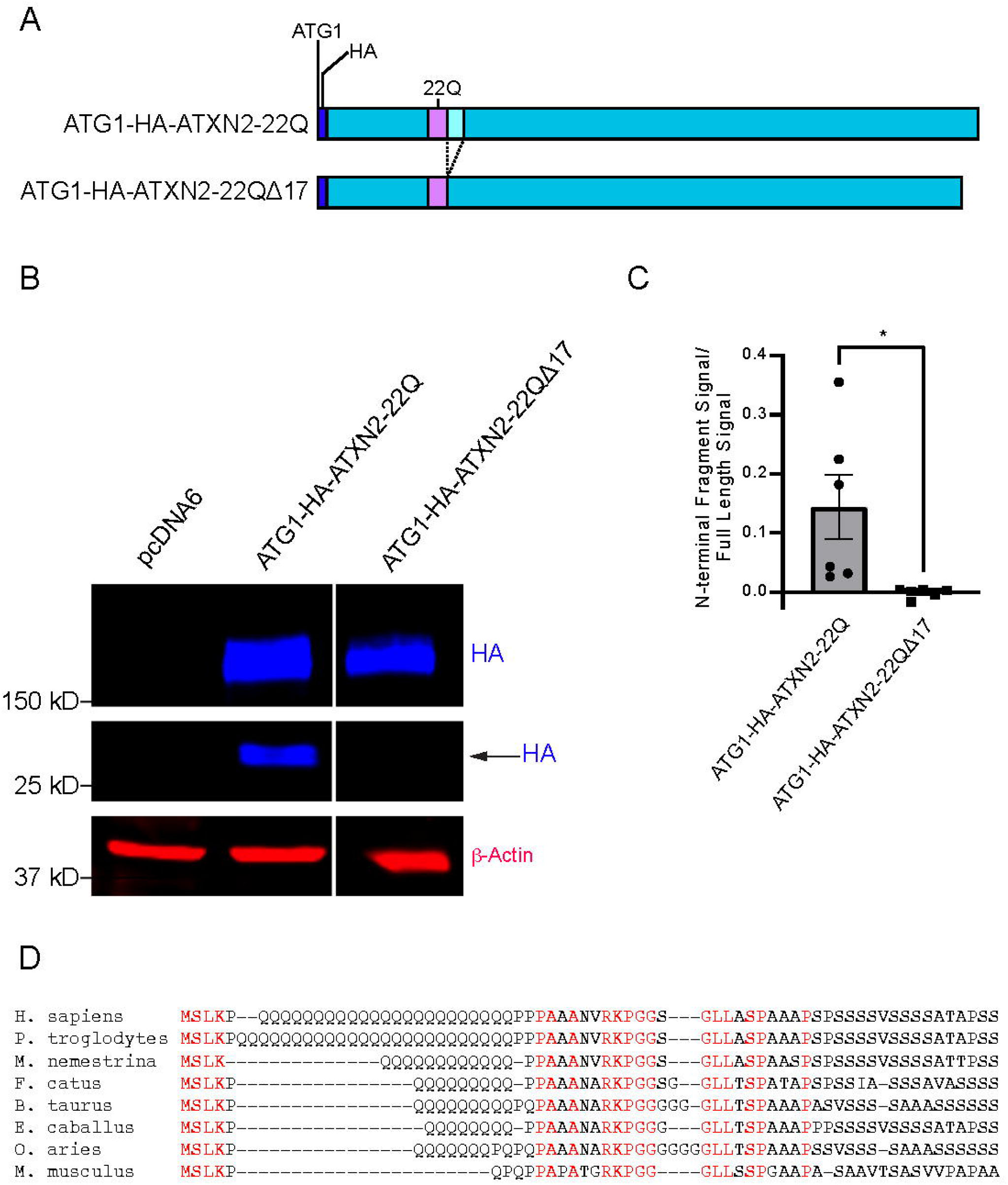
The 17 amino acids downstream of the ATXN2 polyQ are necessary for N-terminal cleavage. **A**. Schematic of the 17 amino acid deletion downstream of the polyQ domain (aqua box). **B**. Probing for the N-terminus of ATXN2 in transfected Hek293E cells via its N-terminal HA tag (HA, blue) shows that ATG1-HA-ATXN2-22Q ⍰17 fails to produce an N-terminal cleavage fragment, while ATG1-HA-ATXN2-22Q does (black arrow). A Western blot representative of six independent experiments is shown. **C.** Quantification of cleavage efficiency. Statistical analysis performed with unpaired T-test. *P < 0.05 **D.** Schematic of amino acid sequence conservation immediately following the polyQ domain in several mammalian species.

To test whether the domain C-terminally adjacent to the polyQ is sufficient for cleavage, we inserted 7 to 90 amino acids of this region between an N-terminal EGFP and a C-terminal mCherry (Fig. 5A). All fusion proteins with 90 to 15 amino acids underwent specific proteolysis that cleaved EGFP from mCherry, while the fusion with only 7 ATXN2 amino acids (residues 188-194) produced very low levels of EGFP fragment at ~27 kDa (Fig. 5B-D). Efficiency of cleavage was statistically significantly reduced when comparing the shortest fusion protein with the two longest, but statistical significance was not reached between the shortest and intermediate length fusions, as overall cleavage efficiency was quite variable across experiments. Importantly however, in all 6 experiments, the fusion protein with only 7 ATXN2 amino acids clearly exhibited the lowest cleavage efficiency (P<10^-5^ to have occurred by chance). These results thus illustrate the importance and sufficiency of the first 17 amino acids following the polyQ domain for ATXN2 cleavage. We also noted that the fusion containing 90 ATXN2 amino acids was cleaved more efficiently than the one containing 15-60 amino acids (Fig. 5C, 5D). Our results thus show that the first 17 amino acids following the polyQ are necessary and sufficient for ATXN2 cleavage, but a broader region might be needed for optimal cleavage efficiency.

**Figure 5.**
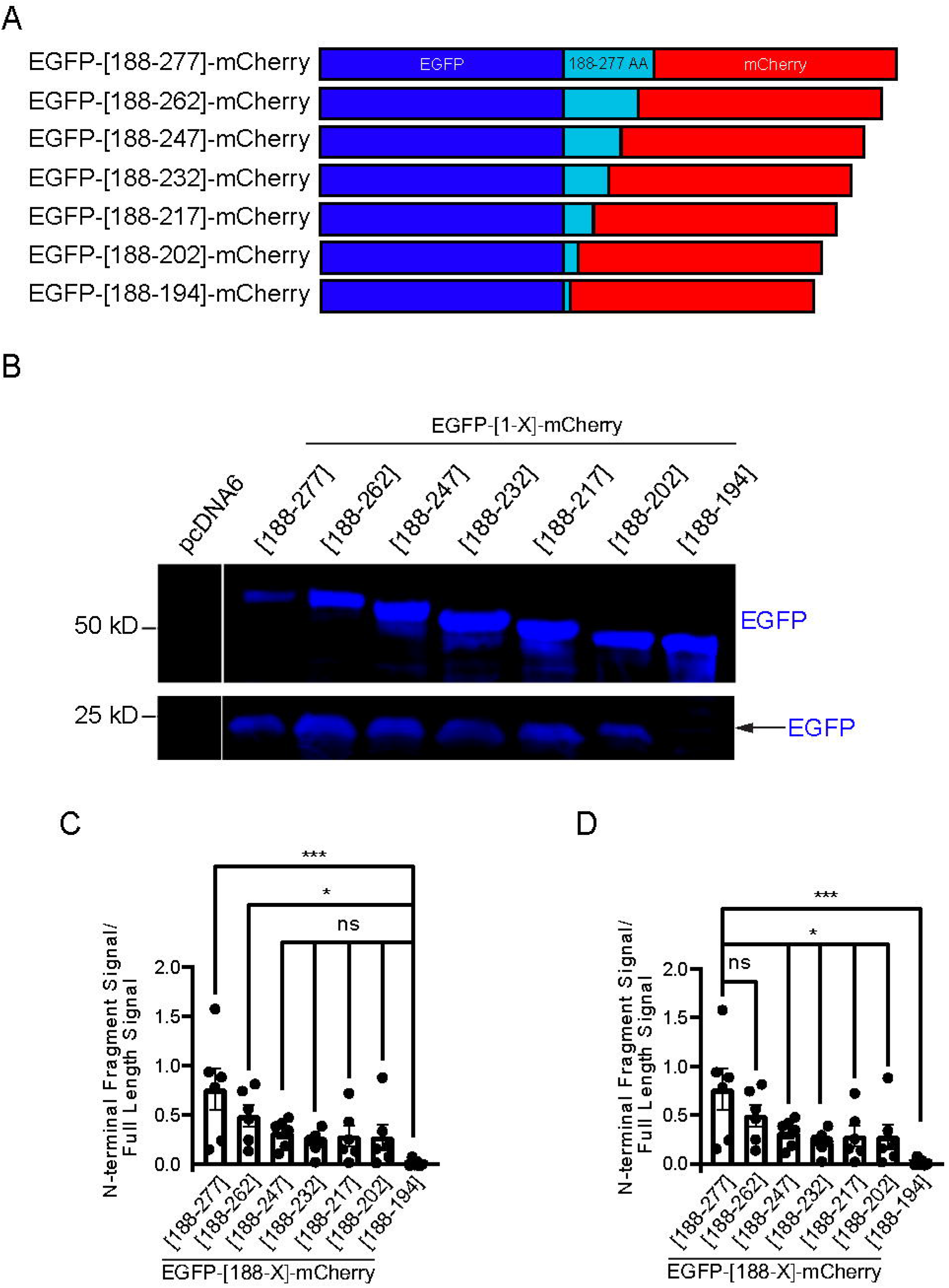
The 17 amino acids downstream of the ATXN2 polyQ are sufficient for proteolytic cleavage to occur. **A**. Schematic of fusion proteins between N-terminal EGFP and C-terminal mCherry moieties with ATXN2 amino acid sequence acting as a linker. 15 amino acids are serially deleted from the ATXN2 sequence in the C to N direction starting from amino acid 277 up to amino acid 194. **B**. Western blot probing for EGFP (blue) from HEK293E lysates expressing these proteins. All fusion proteins were cleaved and produced an N-terminal EGFP fragment except for the fusion protein that contained only 7 ATXN2 amino acids. A Western blot representative of six independent experiments is shown. **C**. Quantification of cleavage efficiency and statistics with EGFP-[188-194]-mCherry as the control group. Statistical analysis performed with one-way ANOVA with Dunnett’s multiple comparisons test. *P < 0.05, ***P < 0.001 **D.** The same quantification and statistics with EGFP-[188-277]-mCherry as the control group. Statistical analysis performed with one-way ANOVA with Dunnett’s multiple comparisons test. *P < 0.05, ***P < 0.001

### ATXN2 produced from the second ATG start codon can also undergo N-terminal proteolysis

The *ATXN2* gene produces multiple mRNA isoforms, but it is usually depicted as encoding a 1313 amino acid protein (starting from ATG1), that includes 165 amino acids N-terminal of the polyQ domain (1, 10, 15, 20, 32–34). The ATXN2 protein is reported to have an apparent molecular weight of about 145 kDa in various humans cell lines and tissue types (4, 5, 10, 20, 33), which would fit closely with a length of 1313 amino acids (predicted MW = 140.3 kD). However, our data show that full-length ATXN2 actually migrates with a ~180 kDa apparent molecular weight, and evidence suggests that the main translational start codon might actually be located just a few codons upstream of those encoding ATXN2’s polyQ domain (19). This second start codon (ATG2) is predicted to produce a significantly shorter protein (predicted MW = 124.4 kD) than the first (ATG1).

To determine whether this shorter ATXN2 isoform also undergoes proteolytic cleavage, we generated HA-tagged constructs that included only ATG2 (Fig. 6A). As with ATG1-containing constructs, HA-tagged ATG2-ATXN2 constructs produced two bands, with the upper band’s mobility dependent on polyQ length, while the faster migrating band was insensitive to polyQ length (Fig. 6B). However, the mobility of these two bands was faster to the mobility of the bands observed with untagged ATG1 constructs, consistent with a shorter protein. While this would be predicted for the slow-migrating band which would correspond to the uncleaved product, this would not be expected if the faster-migrating band corresponded to the C-terminal fragment generated from N-terminal cleavage occurring at the same site as with full-length (ATG1) ATXN2. However, HA staining clearly demonstrated that the slow-migrating band observed with ATG2 contained the N-terminus, while the lower did not (Fig. 6B). Thus, the shorter ATXN2 isoform produced from ATG2 also undergoes N-terminal proteolytic cleavage. We noticed that the difference in size between the two bands detected with C-ATXN2 was smaller than the size difference observed with ATXN2-ATG1 (Fig. 6B), which would suggest that the N-terminal ATG2 fragment should be smaller than the N-terminal ATG1 fragment (probably less than 15 kDa). However, no such faster migrating fragment was observed, even with gels designed for small protein fragments (not shown), though slower migrating bands were observed (Fig. S1C-D). The small N-terminal fragment generated by proteolysis of ATG2-ATXN2 is thus probably unstable.

**Figure 6.**
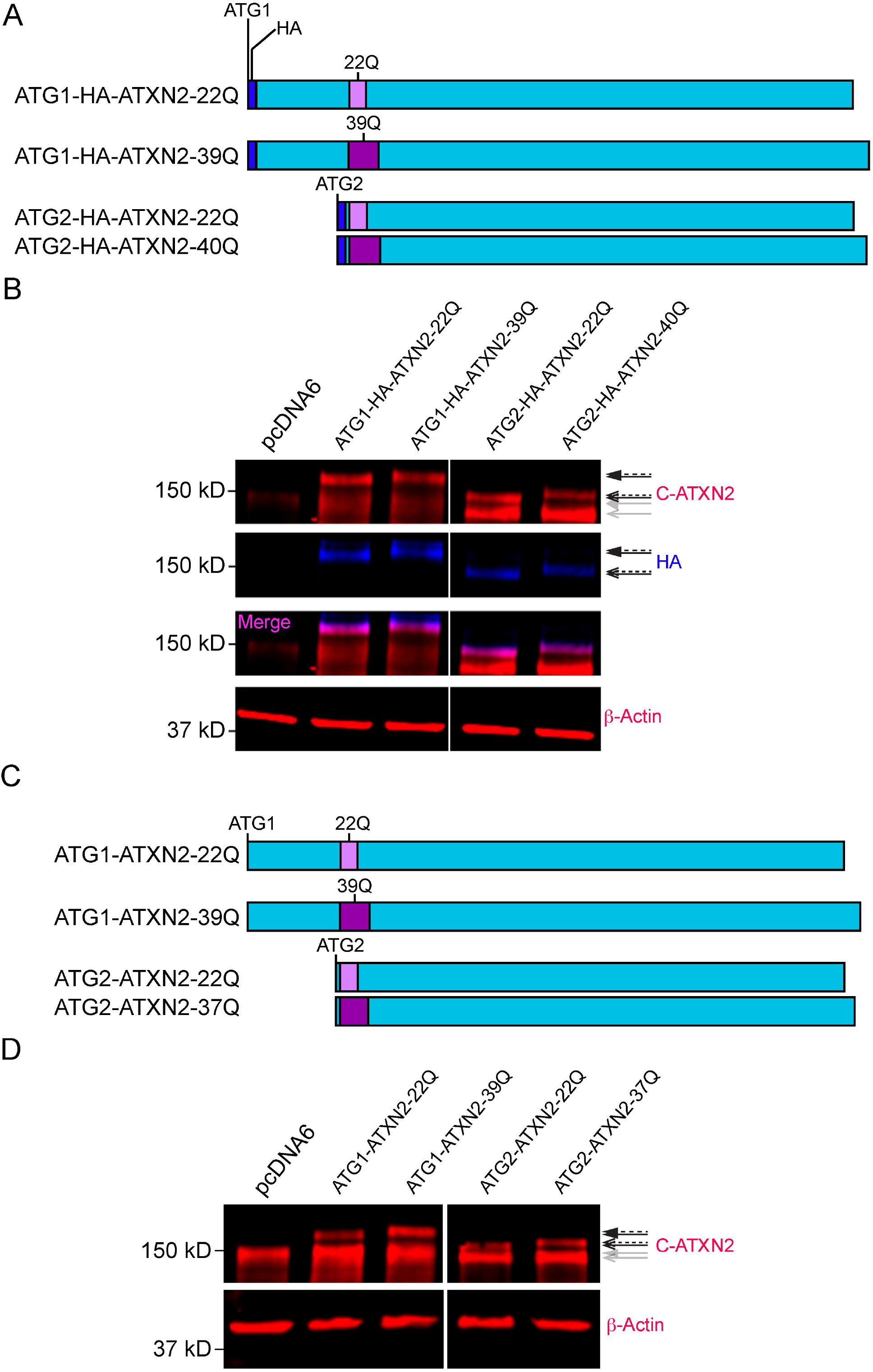
The two ATXN2 isoforms resulting from alternative initiation codon usage undergo N-terminal proteolysis in HEK293E cells. **A**. Schematics of HA-tagged full-length ATXN2 starting at the 1^st^ methionine codon (ATG1, light gray) or at the 2^nd^ known start codon (ATG2, dark gray) that were expressed in HEK293E cells. **B.** Western blot probing with the C-AXTN2 antibody (red) shows doublets suggesting N-terminal cleavage in both ATG1 and ATG2-exoressing lysates. The size migration difference between WT (black flat arrow, solid tail) and mutant (black flat arrow, dashed tail) seen in ATG1 lysates is also seen when comparing WT (black curved arrow, solid tail) and polyQ-expanded (black curved arrow, dashed tail) ATG2 lysates at ~160 kDa. However, the expected ~145 kDa band (gray flat arrow) is instead replaced with a ~140 kDa band in ATG2 lysates (gray curved arrow), suggesting a different cleavage site for the ATG2-ATXN2 isoform. ATG2-ATXN2 behaves similarly to ATG1-ATXN2 when probing for the N-terminus such that only the higher molecular weight bands are detectable via HA (blue), while the C-terminus is present in both bands when probing for C-ATXN2 (red). This is confirmed when overlaying the two channels (magenta). A Western blot representative of three independent experiments is shown. **C.** Schematics for the untagged ATG1 (light gray) versus the ATG2 (dark gray) ATXN2 translation isoforms. **D.** The same migration pattern is seen between ATG1 and ATG2 lysates when probing Western Blots with the C-ATXN2 antibody (red) despite the lack of an N-terminal HA tag, suggesting that the HA tag has no influence on N-terminal proteolysis of ATXN2. A Western blot representative of three independent experiments is shown. Image brightness and contrast were adjusted in both panel B and D for better visualization of full-length (ATG1) ATXN2 proteins, which we were expressed at lower levels than the shorter (ATG2) proteins.

We considered the possibility that the N-terminal HA tag may be influencing N-terminal cleavage behavior, especially when comparing the ATG1 and ATG2 isoforms of ATXN2. ATG1-ATXN2 contains a much longer N-terminal sequence preceding the polyQ domain, while ATG2-ATXN2 only has 5 amino acids in the sequence prior to the polyQ tract. We generated untagged ATG2-ATXN2 constructs with both WT and expanded polyQ domains to compare their cleavage pattern against the untagged ATG1-ATXN2 constructs (Fig. 6C). We found that despite the lack of an HA tag at the N-terminus of ATG1-ATXN2 and ATG2-ATXN2, N-terminal cleavage was conserved in both translation isoforms (Fig. 6D, Fig. S1B and D)). Additionally, removing the HA tag did not change the migration difference seen between the faster migrating bands when comparing ATG1 and ATG2 lysates.

Since the fast-migrating band corresponding to the ATG2-ATXN2 C-terminal cleavage product migrates faster than that of ATG1-ATXN2, we wondered whether it might use a different cleavage-promoting sequence. We therefore deleted the 17 amino acids we found to be necessary for cleavage of full-length (ATG1) ATXN2 from ATG2-HA-ATXN2-22Q (Fig. 7A). The HA signal was detected at ~160 kDa from extracts expressing the protein containing the deletion while the faster migrating band was only detected by the C-terminal antibody, which indicates that when ATXN2 is translated from the ATG2 start site, its cleavage is dependent on a different sequence downstream of the polyQ domain (Fig. 7B). In conclusion, both full-length (ATG1) and short (ATG2) ATXN2 undergo N-terminal proteolytic cleavage in HEK cells, but proteolysis of these two isoforms is likely dependent on different cleavage-promoting sites.

**Figure 7.**
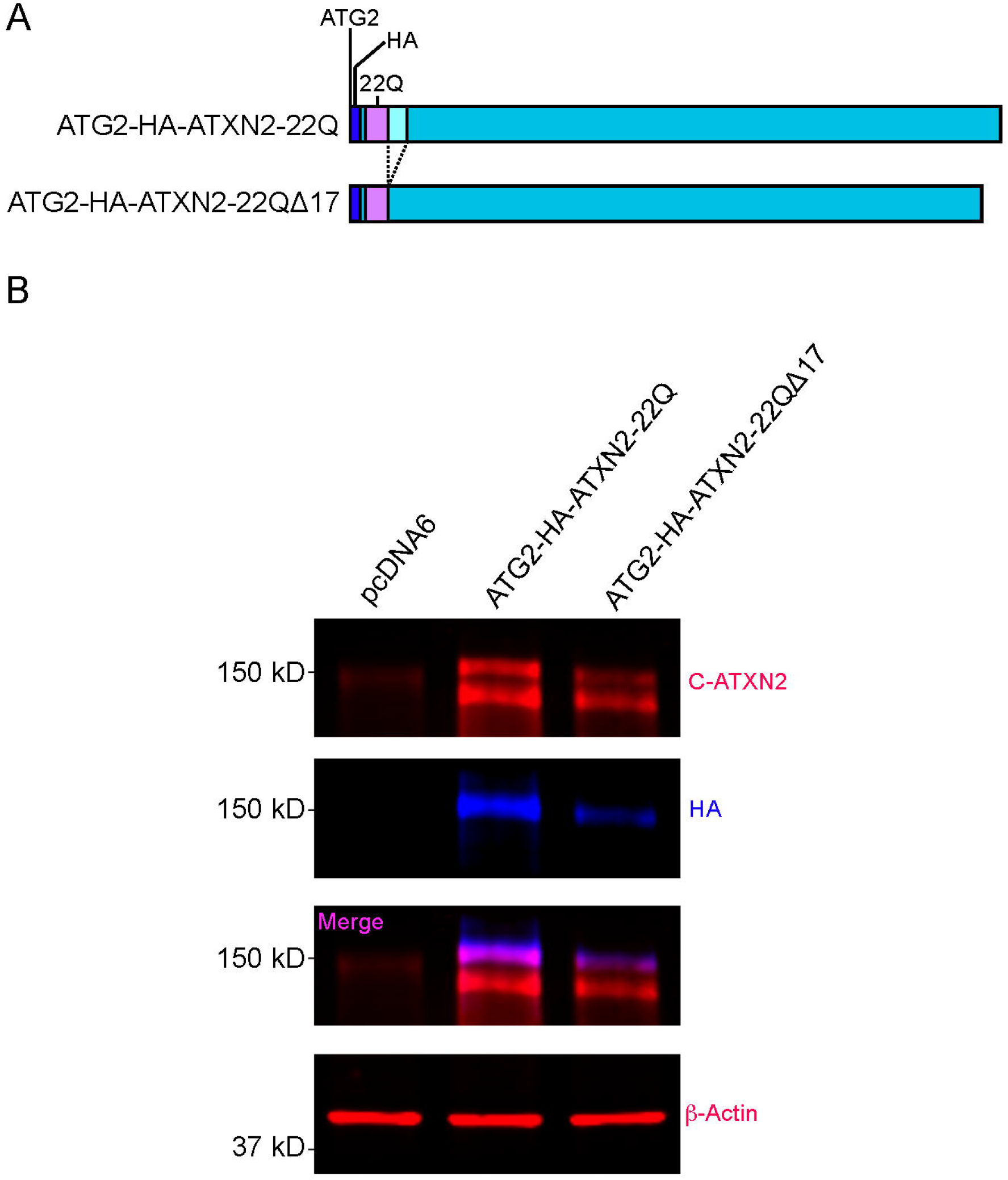
The 17 amino acid sequence downstream of the polyQ domain is not required for cleavage of the shorter ATXN2 translation isoform. **A**. Schematic of the 17 amino acid deletion downstream of polyQ domain from ATG2-HA-ATXN2-22Q sequence (black checkerboard box). **B**. Western blots probed with the HA (blue) and C-ATXN2 (red) antibodies demonstrate N-terminal cleavage for both ATG2-HA-ATXN2-22, and ATG2-HA-ATXN2-22Q ⍰17. A Western blot representative of four independent experiments is shown.

## Discussion

Several polyQ-containing proteins implicated in neurodegeneration are the target of N-terminal proteolysis, which produces small pathological polyQ fragments that accumulate in diseased brains and aggregate (18). Interestingly, polyQ containing ATXN2 fragments have been observed in a brain extract from a deceased patient that carried a 41Q repeat (29). However, N-terminal ATXN2 proteolysis has received little attention since, perhaps because nuclear inclusion body formation with elongated polyQ ATXN2 did not appear necessary for disease progression (29). Our study establishes that when overexpressed in a HEK293E cells, ATXN2 can indeed be targeted by such proteolytic events. Our results are reminiscent of those obtained with the aforementioned SCA2 brain extract (29): as in our study, a small polyQ-containing fragment was observed with the 1C2 antibody that recognizes elongated polyQ stretches, as well as a large, likely full-length protein migrating at about 180kd. The ~145 kDa protein observed in both the patient and a healthy individual was not recognized by the 1C2 antibody. This is however explained by this band corresponding to ATXN2 with a normal polyQ stretch, which is not detected well by 1C2. We note that in a very recent study, a slow-migrating ATXN2 isoform was also observed after transient transfection in Hela, COS7 and HEK293 cells (20), that is most likely identical to the ~180 kDa full-length ATXN2 band we observed, but its nature was not determined in that work. That N-terminal cleavage can be produced in cell culture should be taken into careful consideration by future investigators, particularly when using N-terminal ATXN2 fusion proteins.

Our results show that ATXN2 can be cleaved at a very specific site. Interestingly, our results show that the N-terminus is, itself, a determinant of ATXN2 proteolytic cleavage. Indeed, we identified a 17 amino acid sequence that is necessary and sufficient for proteolysis of full-length ATXN2 generated from ATG1, but this sequence is dispensable for cleavage of the shorter ATG2-dependent ATXN2 isoform. This suggests that ATXN2’s N-terminal conformation and possibly its incorporation into multi-protein complexes might determine how efficiently and where the currently unknown ATXN2 protease is able to cleave ATXN2. It should be acknowledged that while we favor the idea that the 17 amino acid sequence adjacent to the polyQ stretch is the site of ATXN2 cleavage, we cannot exclude that this site is used as a recognition site for the ATXN2 protease that then cleaves ATXN2 at a nearby site.

It is important to note that in fibroblastic cell lines derived from SCA2 patients, or in cell lines in which a long isoform allele was engineered, migration of the main ATXN2 band was observed to be dependent on polyQ length (35). This indicates that in these cells, which do not overexpress ATXN2, this protein is not cleaved N-terminally, or at least is cleaved very inefficiently. It is therefore possible that cleavage is more likely if ATXN2 accumulates excessively or behaves abnormally, because of overexpression or pathology. Clearly, more work is needed to determine *in vivo* in which tissues and under which conditions ATXN2 can be cleaved N-terminally. The functional consequence of the protein cleavage would also be interesting to study. Of the known ATXN2 domains, only the polyQ domain would be missing after cleavage. We would thus predict that the cleaved product is functional, but it might have reduced activity due to lower affinity for its binding partners since polyQ stretches can impact protein interactions (36, 37). This could have important impact on expression of ATXN2 target genes (9).

As mentioned above, the existence of small polyQ-containing N-terminal fragments in at least one SCA2 patient is intriguing (29). Designating ATXN2 as a target of N-terminal proteolysis could open new therapeutic avenues for ATXN2-related neurodegeneration. There are several examples of how either inhibiting N-terminal cleavage of polyQ proteins or targeting their N-terminal cleavage fragments for degradation ameliorate neurodegenerative disease pathology. For example, mutating the caspase 7 cleavage site in ATXN7 produces a cleavage-resistant form of ATXN7, and the disease pathology of SCA7 model mice improves despite an expanded polyQ tract (23). Mice expressing a mutant form of Huntingtin that cannot be cleaved by caspase-6 also fail to develop neurodegenerative phenotypes despite having an expanded-polyQ form of the gene (28). Inhibiting calpain cleavage of ATXN3 improved disease phenotype in SCA3 mice as well (25). Additionally, reducing accumulation of N-terminal cleavage fragments of neurodegenerative polyQ proteins improves phenotypes associated with disease progression in the context of SCA7 (38) and Huntington’s disease (39). In the context of disease, ATXN2 is known to accumulate as shown with N-terminal 1C2 (40, 41) and C-terminal ATXN2 staining (5). Interestingly, cell culture studies have shown that a ~500 amino acid N-terminal fragment of ATXN2 fused to GFP is phosphorylated via Cdk5-p25 and thus targeted for degradation via the proteasome (21). The E3 ubiquitin ligase Parkin might be implicated in this degradation (32). Intriguingly, most putative Cdk5 phosphorylation sites in this ~500 amino acid fragment are located N-terminal of the polyQ. Thus, Cdk5 could be a pharmacological target to promote N-terminal cleavage product clearance. It would therefore be important to determine whether N-terminal fragment accumulation is a factor in the progression of neurodegenerative diseases caused by ATXN2.

## Experimental procedures

### Plasmid Generation, Preparation and Sequence Validation

ATG1-*ATXN2*-22Q and ATG1-*ATXN2*-39Q inserted in the pcDNA6 vector were generously gifted by Dr. Aaron Gitler (Stanford University) (5), and served as starting points for our cloning strategies. To generate novel pcDNA6-based expression constructs, we used either classic cloning approaches with various restriction enzymes and T4 DNA ligase, or generated fragments with overlapping sequences, which were then combined with the NEBuilder Hifi DNA Assembly Kit (New England Biolabs #E5520S). In most cases, we used standard PCR protocols with Taq, Phusion, or Q5 polymerases (New England Biolabs #M0273S,#M0530S, or #M0491S) to generate various vector and insert fragments. However, for CAG-repeat containing fragments, we used a Taq polymerase-based PCR protocol modified from Smith *et al*. (42) that is specifically optimized to amplify GC-rich regions. These PCR reactions contained 10-20 ng of plasmid, 5% DMSO, 1M betaine, 0.10 μM primer mix, and 2.5 mM MgCl_2_. PCR-cycles consisted of 10 minutes of denaturation at 95°C, 9 minutes of annealing at 62°C and a 10-minute extension step.

The coding regions amplified by PCR and inserted in the plasmids were fully sequenced to check for sequence accuracy. For two constructs, we only recovered clones that had a slightly altered CAG repeats. Construct ATG2-*ATXN2*-37Q was missing two CAG repeats. We used this construct since a 37Q stretch is pathological (3, 4, 43). ATG2-*HA-ATXN2*-40Q carried an extra CAG. Plasmids were prepared for transfection using the QIAprep Spin Miniprep Kit (Qiagen #27104). Schematics of the proteins encoded by all the pcDNA6 expression constructs used in this paper are shown in the figures. The coding sequences for the HA-tagged ATG1 and ATG2 ATXN2 proteins (Figs. 1,4, 6, and 7) are localized between the Mlu and XbaI restriction sites of pcDNA6, for the N-terminus-EGFP fusions (Fig. 3) between KpnI and XbaI, and for the EGFP-mCherry fusions (Fig. 5) between KpnI and XbaI.

### HEK293E cells transfection

HEK293E cells were plated in 12-well plates with ~250,000 cells/well. For each transfection except for the various N-terminus-EGFP fusions, 150ng of plasmid was combined with 1.5ml of Mirus-LT1 Transfection Reagent (Mirus Bio #MIR 2300) and 125ml of OptiMEM Reduced Serum Medium (Gibco #31985-070). For the N-terminus-EGFP fusions, the following amounts of plasmid DNA were transfected to adjust protein levels: 150ng of pcDNA6, 300ng of [1-395], 300ng of [1-355], 150ng of [1-315], 50ng of [1-234], and 150ng of [1-194]. After 15 minutes of incubation, each individual transfection mix was added to a single well. Cells were collected and lysed 48-72 hours later.

### Cell lysis and protein extraction from HEK293E cells

Cells were washed with 1mL of ice-cold 1x PBS (Corning) prior to cell lysis. 100ml of ice-cold lysis buffer (Thermo Scientific M-PER Mammalian Protein Extraction Reagent #78501) supplemented with Roche Complete Protease Inhibitor Cocktail (#11697498001) was added to each experimental well, on ice. Plates were shaken at 4°C for 10 minutes. Lysates were spun down at 16000g for 15 minutes at 4°C to remove insoluble material. Protein concentrations were determined using the BCA Protein Assay Kit (Pierce Thermo Scientific #23225). 4x Laemmli Buffer (Bio-Rad Laboratories #1610747) was added and samples boiled at 100°C for 5 minutes.

### Western Blotting

Equal amounts of protein (35-75μg of protein per well depending on experimental condition) were resolved on 10- or 15-well 4-20% Mini-PROTEAN TGX Stain-Free Protein Gels (Bio-Rad Laboratories #4568093 and #4568096) and transferred onto nitrocellulose membranes (BioTrace #66485). Membranes were blocked with either Odyssey Blocking Buffer (TBS) (Licor #927-50000) or soy milk with 20mM Tris-HCL and 0.1% Tween-20. Immunoblotting was performed using the following primary antibodies: mouse anti-C-ATXN2 (BD Biosciences #611378, 1:1000 dilution), rabbit anti-HA (Cell Signaling Technologies #3724S, 1:1000), mouse anti-1C2 (polyQ) (Millipore Sigma #MAB1574, 1:1000), rabbit anti-N-ATXN2 (21^st^ Century Biochemicals, 1:1000), and mouse anti-β-actin (Santa Cruz Biotechnology #sc-47778, 1:10000). Corresponding red and green fluorescent secondary antibodies (Licor #926-68070, #926-32210, and #926-32211, 1:10000) were used for visualization with the Odyssey CLx Infrared Imaging System (LI-COR Biosciences). All Western blot membranes were imaged using the Automatic setting, which acquires images with no saturated pixels without user adjustments. Brightness and contrast of some images were adjusted to better visualize Western blot bands.

### Immunoprecipitation

#### Bead preparation

<u>Protein A</u> Sepharose 4 Fast Flow Beads (GE Healthcare #17-5280-01) were washed with 1mL of ice-cold 1x PBS and then blocked with 0.1% BSA in PBS for 1 hour while rotating at 4°C. Beads were washed twice with 1 mL ice-cold PBS and once with Isotonic Wash Buffer (20 mM Tris-HCl pH 7.5, 150mM NaCl, 0.1% NP-40). Beads were conjugated with the indicated antibodies or immunoglobulin (IgG) at a dilution of 1:125 in 1mL of Isotonic Wash Buffer for 3 hours at 4°C while rotating. Beads were then washed twice with 1mL of ice-cold Isotonic Wash Buffer.

#### Transfection and lysate preparation for immunoprecipitation

HEK293E cells were plated in 100mm^2^ dishes at a density of ~3,000,000 cells/dish and transfected as described above, but with 2.1μg of appropriate cDNA combined with 21ul of Mirus-LT1 Transfection Reagent (Mirus Bio #MIR 2300) and 1.75 mL of OptiMEM Reduced Serum Medium (Gibco #31985-070). 48 hours post-transfection, cells were lysed in 1.0 mL icecold lysis buffer supplemented with Roche protease inhibitor cocktail. Cell lysates were centrifuged at 16,000g at 4°C for 15 minutes and protein concentrations were determined by BCA protein assay. 200μL of lysate (equivalent to ~800 μg of cellular protein) was added to the antibody-conjugated beads and incubated while rotating at 4°C overnight.

#### Immunoprecipitation

Following overnight lysate incubation, beads were spun at 3000g at 4°C for 1 minute. Supernatant was removed and beads were washed 4 times with 1mL ice-cold Isotonic Wash Buffer. Recovered proteins were eluted in 30 μL of 1x Laemmli Buffer (diluted from 2x in cell lysis buffer), 5 minutes at room temperature, followed by a 5-minute incubation at 100 °C. Beads recovered by centrifugation (3000xg, 5 min) and eluates were analyzed by immunoblotting, as described above.

### Western blot quantification and statistical analysis

Quantitative analysis was carried out using ImageStudio Software Version 5.2.5 (LI-COR Biosciences) on non-saturated Western blot bands. The signal intensity of each Western blot band was collected using the Western blot analysis setting on ImageStudio that subtracts the background signal of each lane from the signal of each band of interest. Band signal intensity values were not normalized to a loading control because the values calculated for quantitative analysis were ratios that corresponded to the signal intensity of an individual lane’s cleavage fragment band divided by the signal intensity of its corresponding full-length protein band.

Statistical analysis was performed with Prism 9 (GraphPad). Unpaired t-test was used to determine statistical significance when comparing two sets of measurements. One-way ANOVA followed by Dunnett’s multiple comparisons test was used for experiments with more than two groups. All error bars in graphs represent the standard error of the mean.

## Data availability

All data are contained within the article.

## Acknowledgments

We thank Dr. Aaron Gitler at Stanford University for generously gifting the *ATXN2* plasmids to our lab. We are grateful to the members of the Emery, Anaclet and Weaver labs for fruitful discussions and technical support regarding this project. Furthermore, we thank Drs. David Weaver, Daryl Bosco, and Haley Melikian at UMass Chan Medical School for their careful reading of our manuscript and their helpful comments.

## Funding and additional information

This work was supported by MIRA award 1R35GM118087 from the National Institute of General Medicine Sciences (NIGMS) to P.E.

## Conflict of interest

The authors declare no conflict of interest.

## Abbreviations

ATXN2: Ataxin 2
C-ATXN2: C-terminal ATXN2
HA: human influenza hemagglutinin
PAM2: PABPC1-interacting motif 2
Lsm: like-Sm
Lsm-AD: like-Sm associated domain
EGFP: enhanced GFP

---

**Supplemental Figure 1.** ATG1 and ATG2-ATXN2 protein products in HEK293 cells. **A.** Complete view of a Western blot for ATG1-HA-ATXN2-22Q and ATG1-HA-ATXN2-39Q In addition to the full-length ~180 kDa protein and the ~27 kDa proteolytic product, a ~65-70 kDa band can be clearly and reproducibly detected with the HA antibody (blue). Its migration is sensitive to polyQ length. A band around 115 kDa, insensitive to polyQ length, is detected with C-ATXN2 (red) in both WT and mutant lysates in addition to the major ~145 kDa and ~180 kDa bands. This band could be the C-terminal counterpart of the ~65-70 kDa N-terminal, HA-positive, band. **B.** Western blot for untagged ATG1-ATXN2-22Q and ATG1-ATXN2-39Q. Both WT and mutant lysates show a faint ~115 kDa band (C-ATXN2, red) insensitive to polyQ length. The top half of the membrane was probed with C-ATXN2 antibody, while the bottom half was probed with β-Actin. **C.** Western blot for ATG2-HA-ATXN2-22Q and ATG2-HA-ATXN2-40Q (also shown in Fig. 6B). In addition to the band corresponding to the full-length protein, the former shows two bands at ~30 and ~45 kDa, while the latter show two bands at ~36 and 50 kDa (HA, blue). There is also a band at ~115 kDa in both WT and mutant lysates (C-ATXN2, red) insensitive to polyQ length. The top half of the membrane was probed with C-ATXN2 antibody while the bottom half was probed with β-Actin. **D.** Western blot for untagged ATG2-ATXN2-22Q and ATG2-ATXN2-37Q (also shown in Fig. 6D). Both WT and mutant lysates show a very faint ~115 kDa band (C-ATXN2, red) insensitive to polyQ length. The top half of the membrane was probed with C-ATXN2 antibody while the bottom half was probed with β-Actin.

## Notes

### Competing Interest Statement

The authors have declared no competing interest.

