## Supplementary figures and images for "ATXN2 is a target of N-terminal proteolysis"

### Figure S1

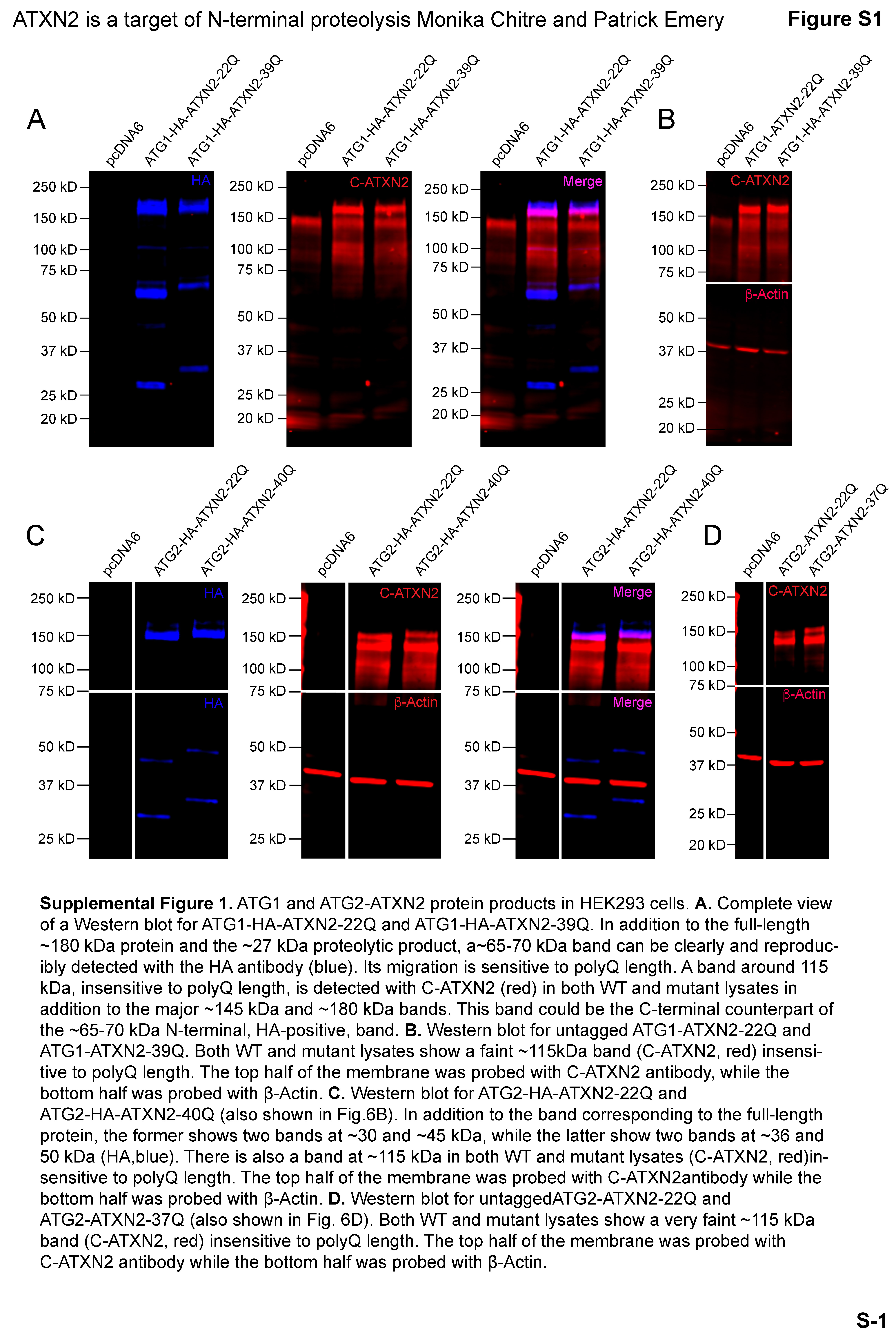
